# Multi-Elemental Isotope And Analog Tracer Application To Measure Nutrient Uptake And Distribution In *Malus* × *Domestica* Borkh

**DOI:** 10.1101/2021.07.20.453131

**Authors:** Raquel Gomez, Lee Kalcsits

## Abstract

Plant nutrient uptake is critical to maintain an optimum balance between vegetative and reproductive growth and fruit quality. Nutrient imbalances, and more specifically the relationship of potassium, magnesium and nitrogen to calcium, can be critical for fruit quality in apple (*Malus* x *domestica* Borkh.). In perennial plants, it is difficult to conduct short-term experiments to measure plant uptake and distribution in response to either cultivar or treatment because of substantial preexisting nutrient pools already present. The use of isotopically labeled nitrogen, and nutrient analogs such as strontium and rubidium allow for the quantification of uptake and distribution that is often not possible with bulk mineral analysis. Here, the objective was to understand how scion genotype influences nutrient uptake and partitioning between aboveground and below ground parts of the tree. In this experiment, 10 atom% ^15^N, Strontium (Sr), and Rubidium (Rb) were applied to three different potted apple cultivars that were either treated with ABA 250 mg/L or 500 mg/L or an untreated control. After 70 days of growth, overall recovery rates of each tracer reflected the mobility of their nutrient analog. Strontium had an average tracer recovery rate of 3.9%, followed by ^15^N with 14.6% recovery and finally Rb with 15.1%. Independent of treatment, ‘Gala’ significantly absorbed more tracer followed by Granny Smith and Honeycrisp for Rb and Sr but not ^15^N. These results have implications in understanding the association between aboveground factors like transpiration and nutrient uptake and distribution in apple.

## INTRODUCTION

Nutrient uptake and distribution by perennial plants is complex because of remobilization and carryover effects of stored nutrient pools. It can be further complicated by the use of composite plants in many horticultural systems where genetically distinct plant parts are used for the below- and aboveground parts of the tree that may produce differences in nutrient uptake and partitioning. Furthermore, in many horticultural crops, elemental balance is critical for ensuring high quality and storability. In apple, the increasingly diverse rootstock and scion genotypes that are available create innumerous combinations that can affect nutrient uptake capacity and scion demand. Although rootstocks have been clearly shown to vary in uptake of individual elements from the soil environment (Fazio et al. 2013; Valverdi et al. 2019) the effects of aboveground factors such as scion genotype and transpiration are less understood. Plant growth and development can affect the nutritional demand. Conversely, nutritional status can also alter growth and development.

Mineral nutrient studies are required to better understand the fundamental process of how plants acquire and allocate mineral elements. These studies can be challenging in perennial plants due to large nutrient pools that exist prior to the start of the experiment and the risk of remobilization of mobile nutrients such as potassium or nitrogen during the course of the experiment. To overcome these challenges, mineral analogs and stable isotopes have been used to track nutrient uptake and distribution to developing plant organs. With the help of mineral analogs like strontium(Sr) (an analog of calcium) (Rosen et al. 2006) rubidium (Rb) an analog of potassium) (Niederholzer and Rosencrance, 2008) and stable isotope tracers like ^15^N (Sturup et al. 2008; Cheng and Raba 2009), uptake and distribution studies can provide more accurate estimates of nutrient uptake under more controlled conditions. These tracers allow for the direct quantification of nutrient movement in response to each treatment without having to account for the large preexisting nutrient pools.

For both potassium and calcium, mineral analogs are better suited tracers because the isotopic forms of both potassium and calcium can be expensive and/or radioactive (Kabata-Pendias 1984). Stable strontium has proven to be an effective analog for calcium uptake because of its equal valence, low mobility in the soil, phloem immobility, plant inability to differentiate between calcium and strontium, and its relatively low abundance in the soil (Kabata-Pendias 1984; Rosen et al. 2006) Similar to strontium, rubidium can also be used as an analog tracer for estimating potassium uptake and transport (McGonicle and Grant 2015). Rubidium has the same valent charge as potassium with similar chemical properties (Niederholzer and Rosencrance, 2008; McGonicle and Grant 2015). Both strontium and rubidium occur naturally in soils in very low concentrations. Therefore, changes in plant rubidium content when supplemental rubidium is added can be used to calculate potassium uptake and distribution (Niederholzer and Rosencrance 2008). The application of ^15^N has been used to trace nitrogen uptake and partitioning in apples (Khemira et al. 1998; Neilsen et al. 2000). As the heavier and less abundant of the two stable nitrogen isotopes, its concentration in plant material can be assessed and quantified when added in excess to the amounts of ^15^N at natural abundance.

Despite the extensive research in plant nutrition, cultivar-specific mechanisms regulating mineral delivery from the roots to aboveground tissues are not well understood. Mineral nutrient mobility plays an integral role in nutrient partitioning. Nitrogen and potassium are highly mobile elements through xylem and phloem pathways, therefore localized deficiencies in the plant stimulates the redistribution of these elements (Zavalloni et al. 2001; Cheng and Raba 2009). However, since calcium is immobile in the phloem, calcium delivery and partitioning is heavily dependent on xylem transport (White and Broadley 2003; de Freitas et al. 2012). Thus, calcium delivery is thought to be highly dependent on water flow through the transpiration stream. For trees with developing fruit, leaves often have greater transpiration than fruit, particularly close to fruit maturity. Therefore calcium delivery is more heavily weighted to leaves (White and Broadley 2003) and concentrations are frequently an order of magnitude difference in concentrations between leaf and fruit tissue.

Previous literature has reported variation in nutrient accumulation amongst plant genotypes (Cheng and Raba 2009; Schloss et al. 2017; Balate et al. 2018). Research has shown some apple scions are better able to accumulate mineral nutrients in both leaves and fruit (de Freitas et al. 2015; Valverdi et al. 2019; Valverdi et al. 2021). For example, a greater accumulation of calcium in ‘Fuji’ when compared to ‘Granny Smith’ has been previously reported (de Freitas et al. 2015). However, these differences in nutrient accumulation by scion type were reported within fruit. Fruit, a major sink organ, has a direct effect on demand and uptake of nutrients. Conversely, it was reported that young non-fruiting ‘Gala’ accumulated more nutrients within its roots, stems and leaves when compared to ‘Honeycrisp’ grown under the same conditions (Valverdi et al. 2019).

In this study, the objective was to measure changes in aboveground mineral nutrient uptake and distribution in non-fruiting apple trees using strontium (analog of calcium), rubidium (analog of potassium) and ^15^N (isotopic tracer of nitrogen) in response to the use of different scion types. The results of this study provide insight into how scion affects mineral nutrient uptake and the distribution between below- and aboveground plant parts.

## MATERIALS AND METHODS

Two year-old *Malus × domestica* Borkh. cv. ‘Granny Smith’, ‘Gala’ and ‘Honeycrisp’ grafted onto M9-T337 dwarfing rootstocks were grown in a pot-in-pot system in soilless potting media (Sunshine mix #2, SunGro Horticulture, Agawam, MA) at Washington State University’s Tree Fruit Research and Extension Center in Wenatchee, WA, with drip irrigation provided four times daily for 30 minutes each. Fertilizer was applied just after budbreak, where 50 g of slow release fertilizer (Osmocote, The Scotts Company, Marysville, OH) and 15 g of granular fertilizer (16-16-16 NPK) was applied to each pot in 2017 and 2018. In 2018, the experiment was set up as a completely randomized design (N = 12) with tracers applied (^15^N, Sr and Rb) or not applied but with balanced fertilization to account for differences from trees where tracers were applied. Tracers were prepared by dissolving 1 g each of each of SrCl_2_ and RbCl, and 200 mg of 10 atom % ^15^NH_4_NO_3_ in a 2 L solution which was evenly poured around the base of the trunk.

### Plant Sampling

After 70 days, trees were destructively separated into leaves, roots and stems. Trees were removed from the soilless media, the roots were washed in distilled water and left to dry, and all leaves were manually removed from the tree. All tissues were dried, and the biomass was recorded. Once dried, the stems were chipped using an electric woodchipper (SunJoe, Carlstadt, NJ), weighed, and ground along with dried leaves to micron size using a VWR high throughput homogenizer (VWR, Radnor, PA).. Roots were ground using a Ninja Master Prep Professional QB1004 (SharkNinja Operating LLC, Needham, Massachusetts). Subsamples of each roughly ground material were sampled from well-mixed media and ground down to micron size using a VWR high throughput homogenizer (VWR, Radnor, PA).

### Nutrient analysis

#### ^15^N-enriched nitrogen isotope analysis

Ground samples were weighed into tin capsules (Costech Analytic Technologies, Inc., Valencia, CA) varying in weight by tissue type: leaf (3 mg), stem (10 mg) and root (3 mg) for nitrogen isotope analysis. Tissue samples of trees receiving ^15^N were sent to the UC Davis Stable Isotope Lab (Davis, CA) using a PDZ Europa ANCA-GSL elemental analyzer interfaced to a PDZ Europa 20-20 isotope ratio mass spectrometer (Sercon Ltd., Cheshire, UK). Samples were interspersed with laboratory standards previously calibrated against NIST standard reference materials. Final delta values were expressed relative to air. Trees not receiving ^15^N were analyzed at the Washington State University Isotope Lab (Pullman, WA using an ECS 4010 Elemental Analyzer (Costech Analytical, Valencia, CA) in conjunction with a Finnigan Delta Plus XP continuous flow isotope ratio mass spectrometer. (Thermo Scientific, Waltham, MA) Samples were interspersed with laboratory standards for calibration. Differences in nitrogen isotope composition were initially expressed as δ^15^N which is calculated as follows

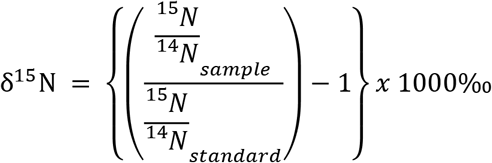

where the relative difference in the ^15^N/^14^N ratio of the sample was compared to the ^15^N/^14^N ratio of a standard (nitrogen gas) and then expressed in permil (‰). at%^15^N was calculated for enriched samplesA by first calculating out the ^15^N/^14^N ratio for each sample using the following equation:

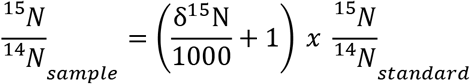

Then at%^15^N for each sample was calculated using:

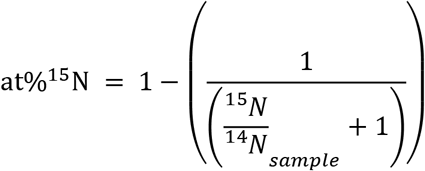

The proportion of ^15^N tracer present in either below- or aboveground plant parts (*i*) was calculated using the following equation:

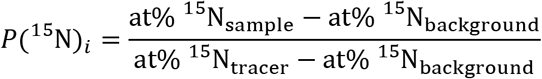

Where a*t%^15^N*_sample_ represents the at%^15^N composition for the below- and aboveground portions of the isotopically labeled plants, at%^15^N_background_ is the ^15^N/^14^N isotopic ratio of samples where no tracer was added, and at%^15^N_tracer_ is the ^15^N/^14^N isotopic ratio of the tracing solution (10 at%^15^N). The amount of ^15^N tracer present into above- and belowground plant parts was then calculated by:

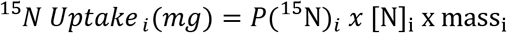

where [N]*_i_* is the N concentration (mg g^−1^) and mass*_i_* is the dry mass of each nitrogen pool (mg).

### Strontium and rubidium analysis

To measure strontium and rubidium tracer uptake, 200±1 mg of ground tissue was weighed into digestion vials and hot-plate digested with 6 mL of HNO_3_. After the digestion was complete, the digest was filtered with a 0.45 μM PTFE filter. (Thermo Fisher Scientific, Waltham, MA) The filtered product was diluted 20x and analyzed using a 4200 MP-AES microwave plasma - atomic emission spectrometer (Agilent Technologies, Santa Clara, CA) and run in combination with elemental ICP standards for validations ((25)). Strontium and rubidium concentrations (mg kg^−1^ dry weight) were calculated using weighted values accounting for concentrations and biomass for below- and aboveground parts. The total uptake for both strontium and rubidium (*i*) into either above- or belowground plant parts (*j*) was calculated using the following equation:

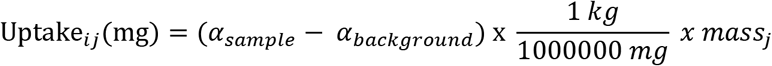

where uptake for below- or aboveground plant parts (*j*) was calculated as the difference between uptake (mg kg^−1^) for the sample where tracer was added (α_i_) and uptake (mg kg^−1^) for the background sample where no tracer was added (α*_background_*).

### Strontium, rubidium, and ^15^N recovery and partitioning

Total plant recovery was calculated from the sum of either ^15^N, strontium, or rubidium (*i*) recovery for all plant parts as follows:

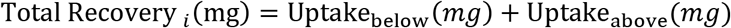

The recovery percentage for ^15^N, strontium, and rubidium in the plant was calculated as:

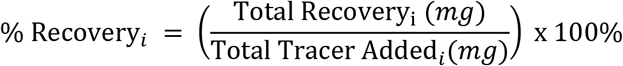

where % Recovery*_i_* represents the total amount of each tracer (*_i_*) taken up into the plant and total tracer added (mg) represents the amount of each tracer element added (73.18 mg, 330.41 mg and 709.22 mg for ^15^N, Sr and Rb, respectively).

The percent partitioned (% Partitioning) to above- or belowground portions (*i*) was then calculated as:

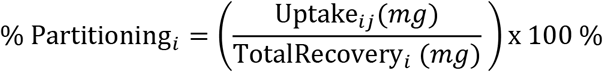

where uptake is the amount of ^15^N, strontium, or rubidium tracer (*i*) recovered (mg) in above- or belowground portions (*j*) of the tree.

### Statistical Analysis

Statistical analysis was done using an analysis of variance in OriginPro 9.1 software (Originlab Corporation, Northampton, MA) and a Q test was performed to eliminate data outliers. Post hoc mean separation was done using Tukey’s honest significant difference test (α = 0.05).

## RESULTS

### Plant Biomass

There were significant differences among cultivars in biomass during the experiment (P=0.017). ‘Granny Smith’ was significantly larger than both ‘Honeycrisp’ and ‘Gala’ (Table 1). ‘Honeycrisp’ had greater root biomass than the other two cultivars (P = 0.058). However, aboveground biomass was significantly greater for ‘Granny Smith’ than both ‘Honeycrisp’ and ‘Gala’ (P = 0.005). Similar to biomass, the root to shoot ratio was not significant affected by treatment. The root to shoot ratio for ‘Honeycrisp’ was greater than 50% but for ‘Granny Smith’, the root to shoot ratio was closer to 40%, with ‘Gala’ intermediate.

**Table 1.** Total biomass distribution (grams dry weight) in above- and belowground portions of the tree for ‘Gala’, ‘Granny Smith’ and ‘Honeycrisp’ apple cultivars (N = 12). Different letters denote mean separations among cultivars using Tukey’s HSD test (α = 0.05).

|  | Aboveground biomass (g) | Belowground biomass (g) |
| --- | --- | --- |
| **Gala** | 353 a | 337 a |
| **Granny Smith** | 433 b | 342 a |
| **Honeycrisp** | 329 a | 357 a |

### Nitrogen

The addition of the tracer into the potting media increased the δ^15^N content of the soil by approximately 40‰. Above- and belowground portions of trees that received ^15^N-enriched nitrogen isotope tracers to the soil were significantly more enriched than trees that received no tracers (P< 0.00001). Of the 73.18 mg of ^15^N applied to each tree, an average of 10.69 mg was recovered, equaling a recovery rate for the ^15^N tracer of approximately 15%. Trees receiving no tracer averaged 0.3662 at%^15^N compared to 0.3863 to 0.3948 for trees receiving ^15^N-enriched nitrogen isotope tracer, for above- and belowground portions, respectively (Table 2). Furthermore, aboveground portions of the tree were significantly less enriched than belowground portions (P = 0.03). Therefore, more ^15^N was present in the aboveground portions of the tree (approximately 60%) compared to only 40% in belowground parts (Figure 1). Furthermore, partitioning of ^15^N was not affected by cultivar. (Figure 1)

**Table 2.** at%^15^N in above- and belowground plant portions for ‘Gala’, ‘Granny Smith’ and ‘Honeycrisp’ apple cultivars (N = 12). Letters denote mean separation among cultivars, compared to a control with no ^15^N added, determined using Tukey’s HSD test (α = 0.05).

|  | Aboveground biomass (g) | Belowground biomass (g) |
| --- | --- | --- |
| **Gala** | 0.3882 a | 0.3920 a |
| **Granny Smith** | 0.3890 a | 0.3929 a |
| **Honeycrisp** | 0.3883 a | 0.3944 a |
| **Control** | 0.3662 b | 0.3662 b |

**Figure 1.**
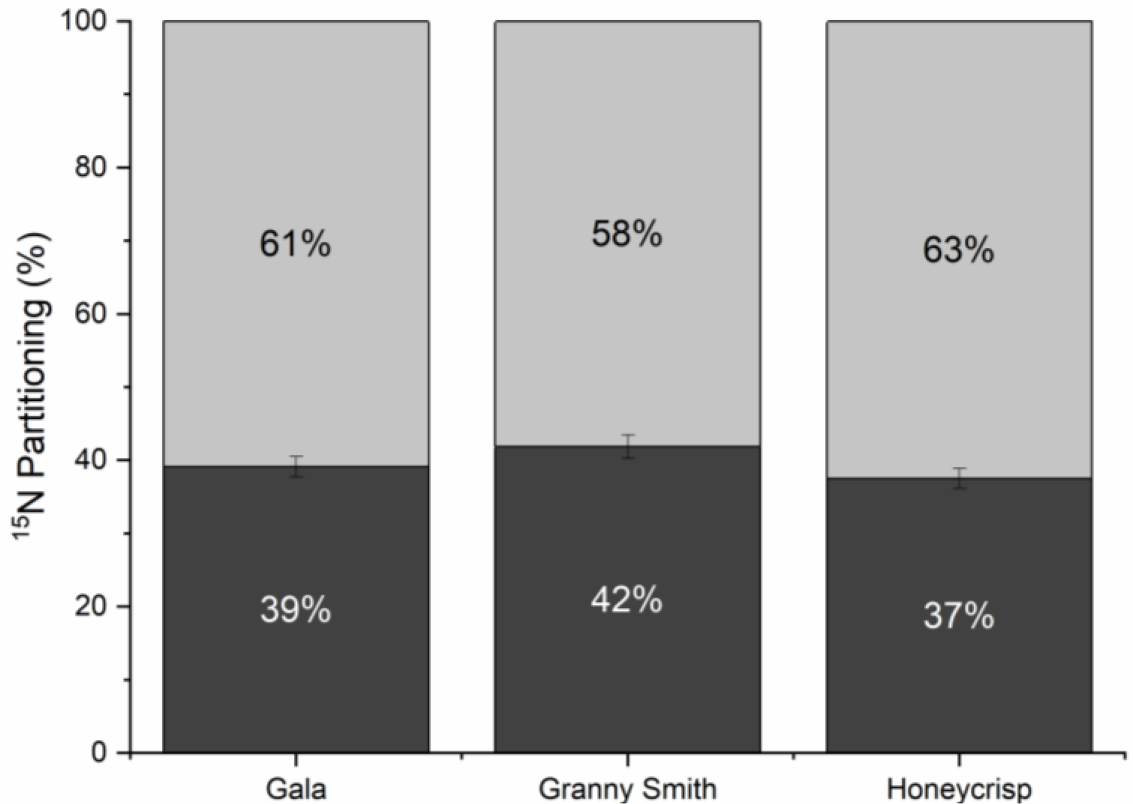
^15^N partitioning between above- (light gray) and belowground (dark gray) plant parts (%) for ‘Gala’, ‘Granny Smith’ and ‘Honeycrisp’ apple trees grown in pots. Error bars denote SEM (N=12).

### Strontium

The natural concentration for strontium was 41.9 mg kg^−1^, and 37.1 mg kg^−1^ for aboveground and belowground portions of the tree, respectively (Table 3). Trees receiving no supplemental strontium had lower strontium concentrations in both above- and belowground sections of the tree (P = 0.002). Of the 330.41 mg of strontium applied to each pot, an average of 12.78 mg was recovered in the tree after 60 days, which is equal to a 3.87% recovery rate. In ‘Honeycrisp’, more strontium tracer was present in the belowground portions (approximately 89%) compared to only 11% in the aboveground portions of the tree (Figure 2). ‘Gala’ accumulated the most strontium when compared to the other two cultivars (Table 4).

**Table 3.** Total strontium present in above- and belowground plant parts (mg kg^−1^) for ‘Gala’ (GA), ‘Granny Smith’ (GS) and ‘Honeycrisp’ (HC) apple cultivars (N = 12). Letters denote mean separations among cultivars determined using Tukey’s HSD test (α = 0.05).

|  | Aboveground biomass (g) | Belowground biomass (g) |
| --- | --- | --- |
| **Gala** | 70.7 c | 76.4 c |
| **Granny Smith** | 46.2 ab | 45.8 ab |
| **Honeycrisp** | 52.0 b | 55.8 b |
| **Control** | 41.9 a | 37.1 a |

**Table 4.**
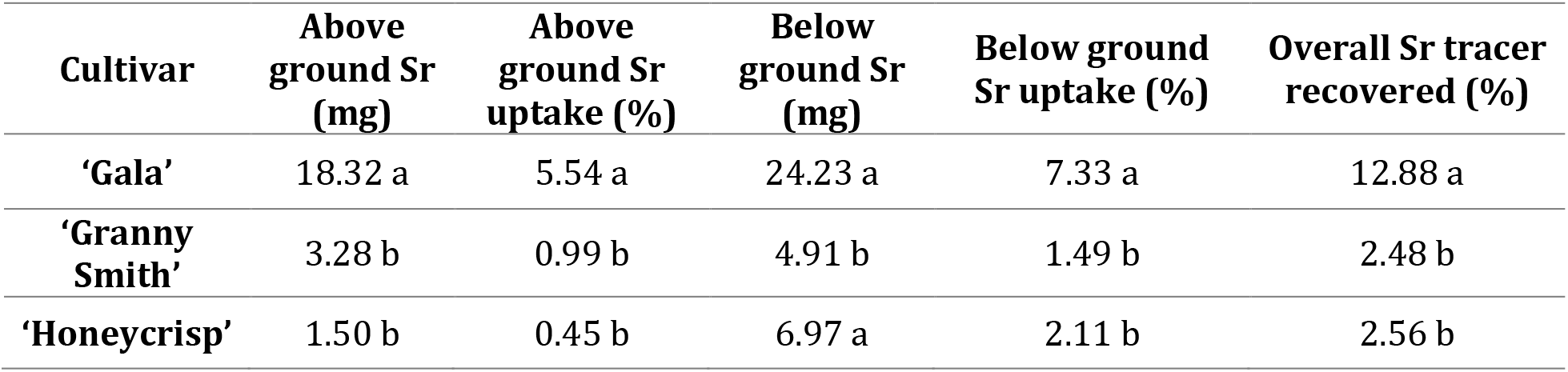
Total strontium present in above and belowground plant parts (mg) and total strontium uptake (%) as well as total strontium tracer recovery in above- and belowground plant parts for ‘Gala’ (GA), ‘Granny Smith’ (GS) and ‘Honeycrisp’ (HC) apple cultivars (N = 12). Letters denote mean separations among cultivars determined using Tukey’s HSD test (α = 0.05).

**Figure 2.**
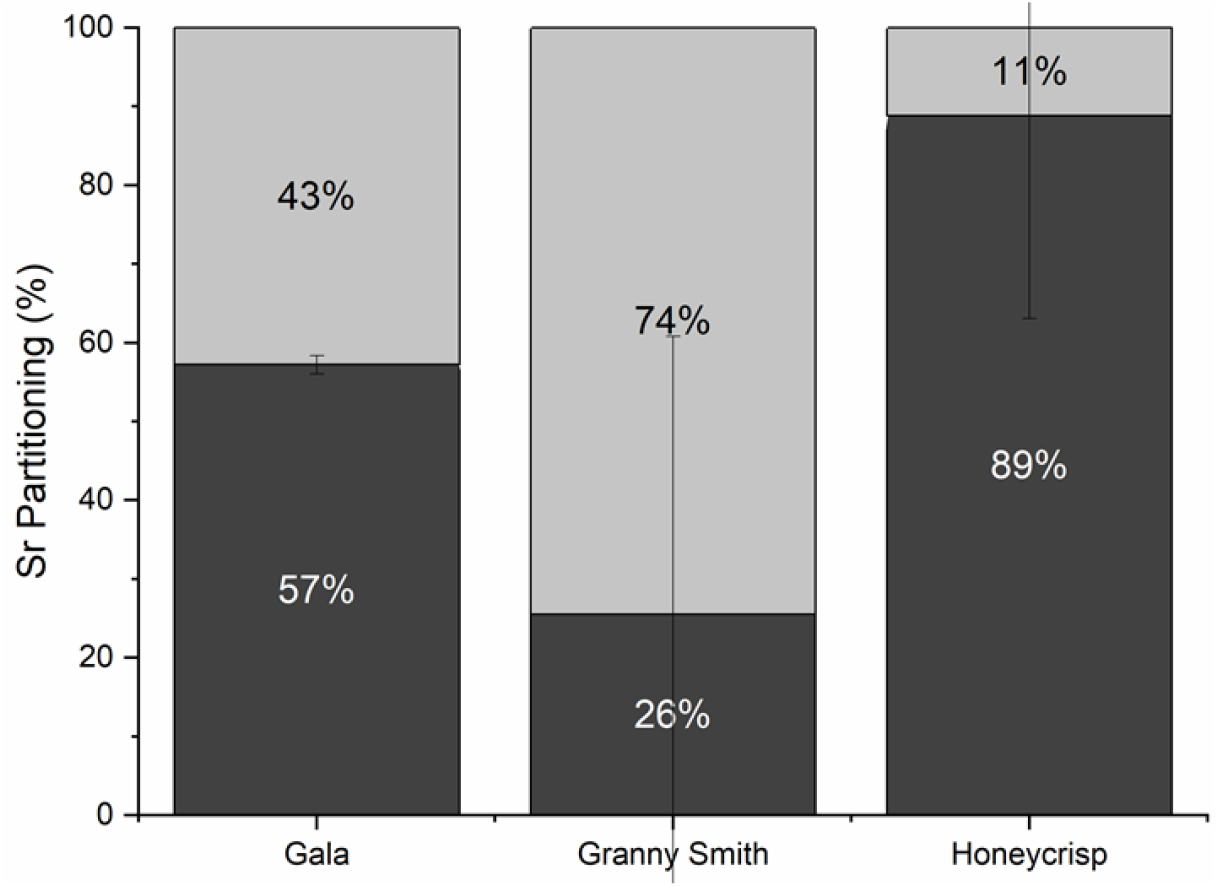
Strontium partitioning between above- (light gray) and belowground (dark gray) plant parts (%) for ‘Gala’, ‘Granny Smith’ and ‘Honeycrisp’ apple trees grown in pots. Error bars denote SEM (N=12).

### Rubidium

Natural rubidium concentrations were 13.7 mg kg^−1^ and 19.1 mg kg^−1^ in above- and belowground sections of the trees, respectively (Table 5). Trees not receiving supplemental rubidium had significantly lower tissue concentrations compared to trees that had received supplemental rubidium (Table 5). Of the 709 mg of supplemental rubidium applied, an average of 107 mg was recovered, which amounted to 15.1% recovery rate. However, unlike strontium, no significant differences among cultivars or treatments were observed (P > 0.05). Almost three quarters of the rubidium tracer detected in the trees was present in the aboveground sections (Figure 3).

**Table 5.** Total rubidium present in above- and belowground plant parts (mg kg^−1^) for ‘Gala’ (GA), ‘Granny Smith’ (GS) and ‘Honeycrisp’ (HC) apple (N = 12). Letters denote mean separations among cultivars determined using Tukey’s HSD test (α = 0.05).

|  | **Aboveground biomass (g)** | **Belowground biomass (g)** |
| --- | --- | --- |
| **Gala** | 286.7 a | 127.1 a |
| **Granny Smith** | 184.1 a | 75.6 a |
| **Honeycrisp** | 233.4 a | 96.8 a |
| **Control** | 13.7 b | 19.1 b |

**Figure 3.**
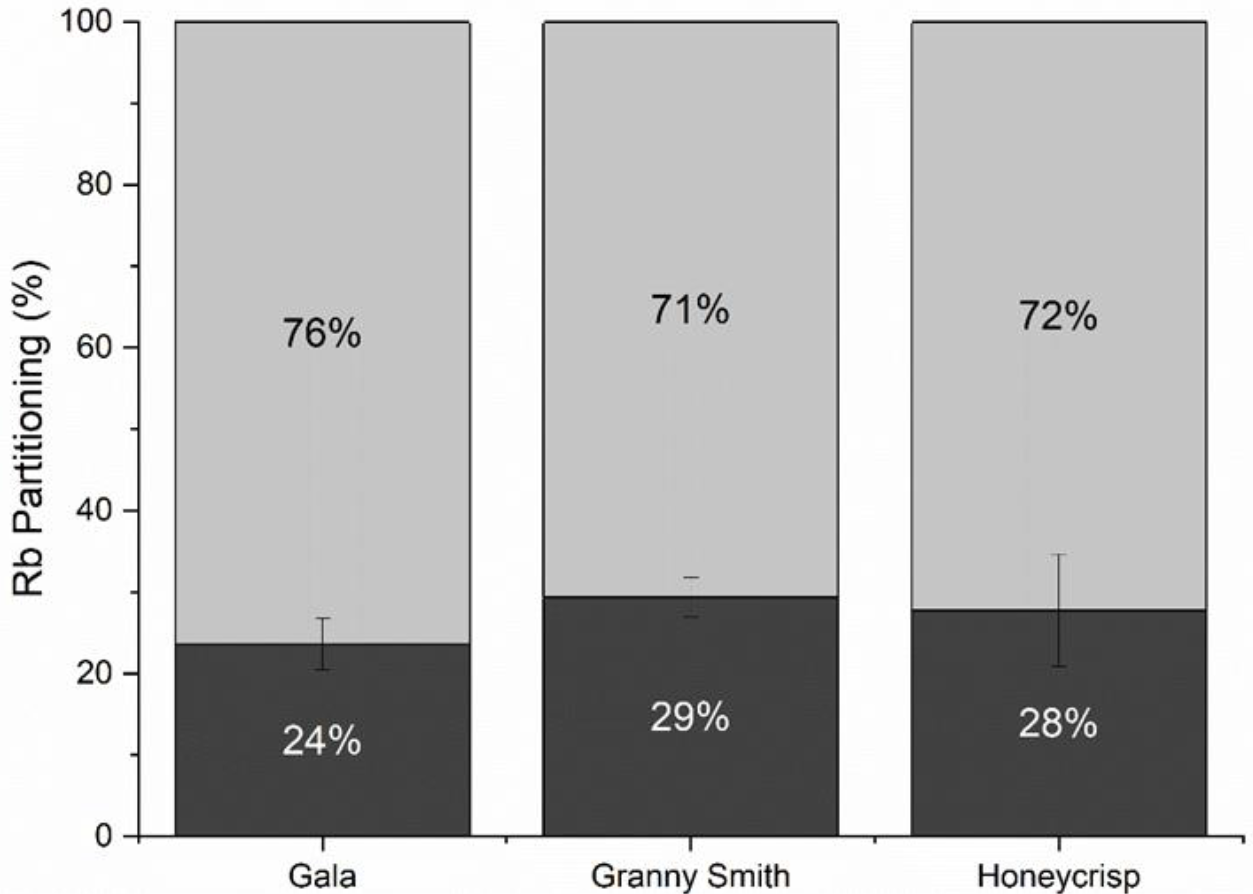
Rubidium partitioning between above- (light gray) and belowground (dark gray) plant parts (%) for ‘Gala’, ‘Granny Smith’ and ‘Honeycrisp’ apple trees grown in pots. Error bars denote SEM (N=12).

## DISCUSSION

Root uptake of strontium was affected by aboveground factors such as scion genotype and transpiration through the application of exogenously applied ABA but rubidium and nitrogen uptake were not. Strontium uptake was lower than both rubidium and nitrogen and was less preferentially allocated to aboveground tissue.

Here, using rubidium as a potassium analog, strontium as a calcium analog, and ^15^N as a nitrogen tracer, we report how nutrient uptake and partitioning was different among elements and highlight the importance of internal and external aboveground factors influencing nutrient uptake and distribution. Radioisotopes, stable isotopes and mineral analogs have been used as tracers to study nutrient uptake and distribution within plants. Plants do not differentiate rubidium from potassium (Menzel 1954; Relman 1956; Lauchli and Epstein 1970; Drobner and Tyler 1998). Therefore, rubidium can be used as an analog for potassium uptake and movement within plants. In this experiment, rubidium was used as an analog tracer for potassium mobility and partitioning with the plant. Strontium has been used as a tracer in a similar role (Rosen et al. 2006) as an analog for calcium. ^15^N has been used extensively to trace nitrogen movement in plants. Previous research has reported much higher recovery raters than those observed here (14.6%), reaching 50.5% ^15^N recovery (Morris et al. 2018) as well as the 20-30% recovery reported in an earlier studies (Khemira et al. 1998; San-Martino et al. 2010). One of the major factors potentially influencing the low recovery rate is the short window of tracer application to final destruction (70 days) as well as the timing of tracer application. The studies mentioned look at nutrient uptake and accumulation over a full growing season from bloom to apple harvest (Neilsen et al. 2000). Therefore, recovery rates observed here are in line with expected tracer accumulation during the experimental period.

### Scion affected overall uptake and partitioning for strontium but not nitrogen or rubidium

Scion genotype affected strontium uptake but not nitrogen or rubidium. ‘Gala’ had the greatest strontium recovery at 7.02%, followed by ‘Honeycrisp’ at 3.08% and ‘Granny Smith’ at 1.36%. (Figure 2). Previous research has reported genotypic variation in nutritional demand. For example, scion-specific calcium demand has been reported throughout the growing season where ‘Gala’ was reported to be a moderate calcium accumulator (Zheng et al 2006). Previous work reported low calcium concentrations and high potassium and nitrogen concentrations and consequently, high susceptibility to mineral imbalance disorders for ‘Granny Smith’ but not ‘Fuji’ apple fruits (de Freitas et al 2015). ‘Gala’ historically has shown limited bitter pit development (Ferguson and Watkins 1989). While scion variation has been linked to unbalanced allocation of elements to developing fruit or at the cellular level (de Freitas et al 2012; de Freitas et al 2015), the differences observed here were related to uptake rather than distribution within the scion. This may indicate a possible feedback between the scion and the rootstock that regulates nutrient uptake and transport to aboveground tissues. This scion to rootstock regulation has been previously identified in many species like mangoes, tomatoes and cotton (Schmutz and Ludders 1999; Chen et al. 2003; Wang et al. 2012). Previous research has shown apple scion genotypes can affect biomass partitioning (Tworkoski and Fazio 2016) but research has not previously shown a scion mediated effect on nutrient demand and distribution.

Rubidium uptake and distribution to the scion was the greatest for ‘Gala’ when compared to the other two cultivars. (Figure 3) Although overall uptake rates differed among rubidium and strontium, these results were similar to those observed for strontium. This may underscore the effect of scion on whole plant nutrient acquisition and distribution. It is possible that ‘Granny Smith’ trees, which were the most vigorous scion genotype, had greater nutrient demands in the aboveground portions of the trees due to their greater leaf area and biomass. Rubidium accumulation was lower in aboveground tissue for both ‘Honeycrisp’ and ‘Granny Smith’ indicating that even for mobile nutrients like rubidium, or potassium, the transpiration demand of leaves can still influence overall demand.

## CONCLUSION

Here, we demonstrated the use of isotope and mineral analog tracers as tools to better understand the effect of aboveground factors such as scion genotype on nutrient uptake and distribution within potted apple trees. Strontium, as a calcium analog, was taken up and allocated to the shoot more slowly than nitrogen and rubidium, a potassium analog. Scion genotype had a significant effect on strontium uptake with ‘Gala’ accumulating more of each tracer than either ‘Granny Smith’ or ‘Honeycrisp’. More research is needed to discern the underlying mechanisms contributing to these genotypic differences and selectivity in nutrient uptake and partitioning. In addition to previous research showing scion genotype effects on nutrient partitioning between fruit and leaves and cellular allocation within the fruit, there is a significant contribution of aboveground factors that affect overall nutrient uptake and partitioning in apple that needs to be more closely considered in future research and management decisions.

## ACKNOWLEDGMENTS

1014919. The authors would like to acknowledge Hector Camargo and Katie Mullin for their technical assistance and Chris Sater for reviewing the manuscript.

